# Understanding Antibody-Target Antigen Interactions and the Avidity Effect Using Mathematical Modelling

**DOI:** 10.1101/2024.05.10.593537

**Authors:** Luke A. Heirene, Helen M. Byrne, Eamonn A. Gaffney, James W. T. Yates

## Abstract

**Background and Purpose:** Immunotherapies are designed to exploit the immune system to target pathologies, for instance, but not exclusively, cancer. Monoclonal antibodies (mAbs) are an important class of immunotherapies that induce anti-tumour effects in numerous ways. Fundamental to the success of mAbs in cancer treatments are their interactions with target antigens. For example, binding multiple antigens, which increases binding affinity, termed the avidity effect, has been shown to impact treatment outcomes. However, there has been limited theoretical analysis addressing the impacts of antibody-antigen interactions on avidity, potency, and efficacy. Hence, our aim is to develop a mathematical model to develop insight on these impacts.

**Experimental Approach:** We analyse an ordinary differential equation model of bivalent, monospecific IgG antibodies binding to membrane bound antigens. We obtain equilibrium solutions of the model and conduct a global parameter sensitivity analysis to identify which antibody-antigen interactions impact quantities, such as antigen occupancy, that contribute to mAb potency and efficacy.

**Key Results:** We show that the ratio of antibody to antigen number impacts antigen occupancy, bound antibody number and whether an antibody can bind both its antigen-binding arms. A sensitivity analysis reveals that antigen occupancy and the ratio of bound antibody to total antigen number are sensitive to the antibody-antigen binding rates only for high antibody concentrations. We identify parameter ranges in which the avidity effect is predicted to be large for antigen occupancy and bound antibody numbers.

**Conclusion and Implications:** These results could be used in the preclinical development of mAb therapies by predicting conditions which enhance mAb potency, efficacy, and the avidity effect.

## 1 Introduction

The immune system can target infection and diseases such as cancer (Gonzalez et al. 2018). It comprises the innate system, which responds to a broad range of threats within the body, and the adaptive system, which is highly specific to its target (Chaplin 2010). The innate and adaptive systems work in unison to prevent the development of cancer and to destroy any cancer cells that arise (Liu and Zeng 2012).

However, tumour cells can disrupt the immune response in a variety of ways. For example, they can inhibit the immune response by expressing inhibitory immune checkpoint receptors such as the programmed cell death-ligand 1 (PD-L1) (Vinay et al. 2015). One aim of cancer immunotherapies is to mitigate the ability of tumour cells to disrupt the immune system while another is directly enabling the immune system to target the cancer.

Anti-tumour monoclonal antibodies (mAbs) are a class of immunotherapies that have had major clinical impact via both mechanisms (Rajewsky 2019). For example, mAbs can induce anti-tumour effects by binding to immune checkpoint receptors on tumour cells and, in so doing, inhibiting the ability of tumour cells to suppress the immune response (Haanen and Robert 2015). MAbs can also induce their anti-tumour effects through antibody effector functions where they recruit and stimulate other parts of the immune system to target and kill cancer cells (Erp et al. 2019). An important antibody effector function is antibody-dependent cellular cytotoxicity (ADCC), where mAbs that bind to cancer cells enable immune cells to recognise and then actively lyse the cancer cells (Nigro et al. 2019).

Regardless of how mAbs induce their anti-tumour effects, central to their potency and efficacy are their interactions with target antigens,(where potency is defined as the expression of drug activity for a given concentration or the amount of drug required to produce a defined effect, while efficacy is defined as the maximum response that can be achieved regardless of dose). Mazor, Yang, et al. 2016 highlight the relationship between antibody-antigen interactions and effector function potency and efficacy. By modifying the affinity of a variety of mAbs for their target antigen, Mazor, Yang, et al. 2016 showed that lower affinity variants exhibited improved effector function. Mazor, Yang, et al. 2016 hypothesised that lower affinity variants were more likely than higher affinity variants to form monovalent than bivalent bonds. Increased levels of monovalent binding increase the number of antibodies per target antigen that can activate an immune cell to kill a tumour cell and, thereby, increase both effector function potency and efficacy. Increased target antigen occupancy has also been found to enhance immune checkpoint inhibitors (Junker et al. 2021).

In separate experimental work, Bondza et al. 2020 showed how the proportion of monovalently and bivalently bound antibodies change as antibody concentration varies. For most concentrations, antibodies are predominantly bivalently bound but for very high concentrations, they are mostly monovalently bound. A schematic of the key antibody-target interactions detailed in Mazor, Yang, et al. 2016 and Bondza et al. 2020 is presented in Figure 1. Together, the works of Mazor, Yang, et al. 2016 and Bondza et al. 2020 show how antibody-antigen interactions on the surface of the target cell can alter the antibody’s binding state and impact mAb potency and efficacy. However, there is no consensus about which of the parameters governing antibody-antigen interactions are most important for mAb potency and efficacy. For example, is it better to have a strong binding affinity or a larger number of target antigens? How does this change as the antibody concentration varies? As a first step towards answering these questions, in this paper we use mathematical modelling to investigate how factors that regulate antibody-antigen interactions (e.g. on and off binding rates, antibody concentration and antigen density) impact the potency and efficacy of mAb therapies, as quantified for instance by antigen occupancy and the number of bound antibodies.

**Fig. 1:**
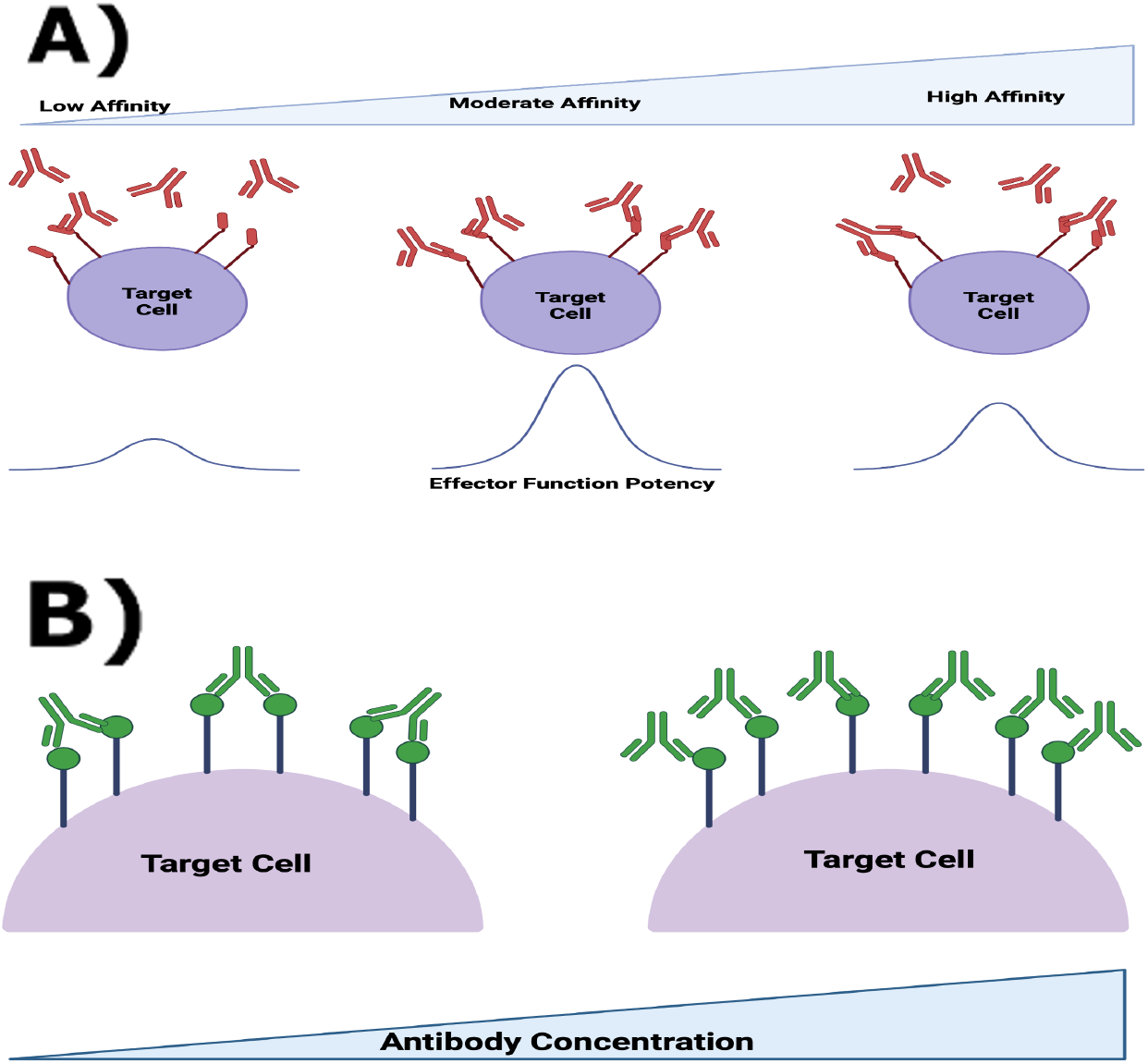
Schematic detailing the key antibody-antigen interactions involved in mAb therapies. A): Enhancing effector function potency by modifying the antibody’s affinity for its target antigen (adapted from Mazor, Yang, et al. 2016). Effector function potency is improved for antibodies with moderate affinity due to the increased number of antibodies bound to the cell. The increased number of antibodies bound to the cell is due to increased levels of monovalent binding. Antibodies with high affinity are more likely to be bivalently bound and, therefore, the total number of antibodies bound to the cell is reduced. The curves depicted are a representation of the resulting potency from different affinity mAb variants. B): The binding state of the antibody changes with antibody concentration (Bondza et al. 2020). For low antibody concentrations, antibodies are primarily bivalently bound. As the antibody concentration increases, antibodies are primarily monovalently bound. Created with Biorender.com.

When developing a mAb therapy, a key consideration is how the number of antigens an antibody can bind, termed its valency, affects its target binding dynamics. Consider, for example, a monospecific Immunoglobulin G (IgG) antibody, a common type of bivalent antibody. After binding to a target antigen, the antibody can bind to a second target antigen with its remaining binding arm. Binding multiple antigens increases the experimentally measured binding affinity, termed the “avidity” effect (Oostindie et al. 2022). Some cells infected with pathogens, such as a virus, can evade an avidity effect if the surface proteins targeted by the antibodies are immobile and sparsely distributed across the cell membrane. In such cases, it is rare for an antibody to bind both of its arms, resulting in lower overall binding affinity and, hence, smaller levels of antibody binding at non-saturating antibody concentrations (Klein and Bjorkman 2010). This is just one example where avidity, or lack thereof, is known to contribute to antibody therapeutic effect. For other examples, see the review by Oostindie et al. 2022.

Clearly, the avidity effect is an important feature of antibody-antigen interactions. However, there is no consensus about the conditions under which the avidity effect is large for mAb binding. Mathematical modelling can be used to investigate whether there are certain ranges of total target antigen densities and binding affinities that result in a large avidity effect.

Several authors have developed mathematical models that describe antibody binding to cells (Perelson and DeLisi 1980; Kaufman and Jain 1992; Rhoden et al. 2016; Sengers et al. 2016). These models differ in the mAb-antigen interaction under consideration (e.g monospecific or bispecific) and the way in which the second arm of the antibody binds to its target antigen. For example, Sengers et al. 2016 assumes that binding of the second arm is limited by surface diffusion of target antigens whereas Rhoden et al. 2016 assumes that binding of the second arm is driven by antigen levels within reach of the bound arm of the antibody.

Here, we focus on monospecifc, bivalent antibodies. We will extend the model of bivalent ligand-antigen binding given in Perelson and DeLisi 1980 with two main aims. First, we aim to investigate how simultaneously varying system parameters, such as antigen density and binding affinity, impacts quantities of interest such as the antibody binding state, the number of bound antibodies and antigen occupancy. Secondly, we aim to identify parameter ranges which induce a strong avidity effect.

## 2 Methods

In this section, we introduce a time-dependent mathematical model that describes the binding of a monospecific, bivalent antibody to target antigens on the cell membrane. We formulate our model as a system of ordinary differential equations (ODEs).

Our model is based on an existing model of bivalent ligand binding presented in Perelson and DeLisi 1980. Unlike Perelson and DeLisi 1980, we do not assume that ligand (antibody in this case) is always in excess. We formulate our model in units of the number of antibody or antigen rather than concentration in order to more clearly calculate quantities of interest like antigen occupancy and the ratio of bound antibody to antigen. As in Perelson and DeLisi 1980, we neglect spatial effects by assuming that the system is well mixed and that target antigens are distributed uniformly over the cell membrane. The dependent variables are the number of unbound target antigens, *r*(*t*); the number of unbound antibodies, *A*_0_(*t*); the number of monovalently bound antibodies, *A*_1_(*t*); and the number of bivalently bound antibodies, *A*_2_(*t*).

The mathematical model we consider describes how a monospecific, bivalent antibody binds to a target antigen on a single tumour cell as depicted in Figure 2. We assume an unbound monospecific antibody binds reversibly with one of its arms to a free target antigen, to form a monovalently bound antibody-antigen complex. The monovalently bound antibody reversibly binds a second free antigen with its unbound arm, to form a bivalently bound antibody. At this stage, each antigen-bound arm may dissociate from its antigen. The rate at which one arm dissociates is assumed to be equal to, and independent of, the rate at which the other arm dissociates.

**Fig. 2:**
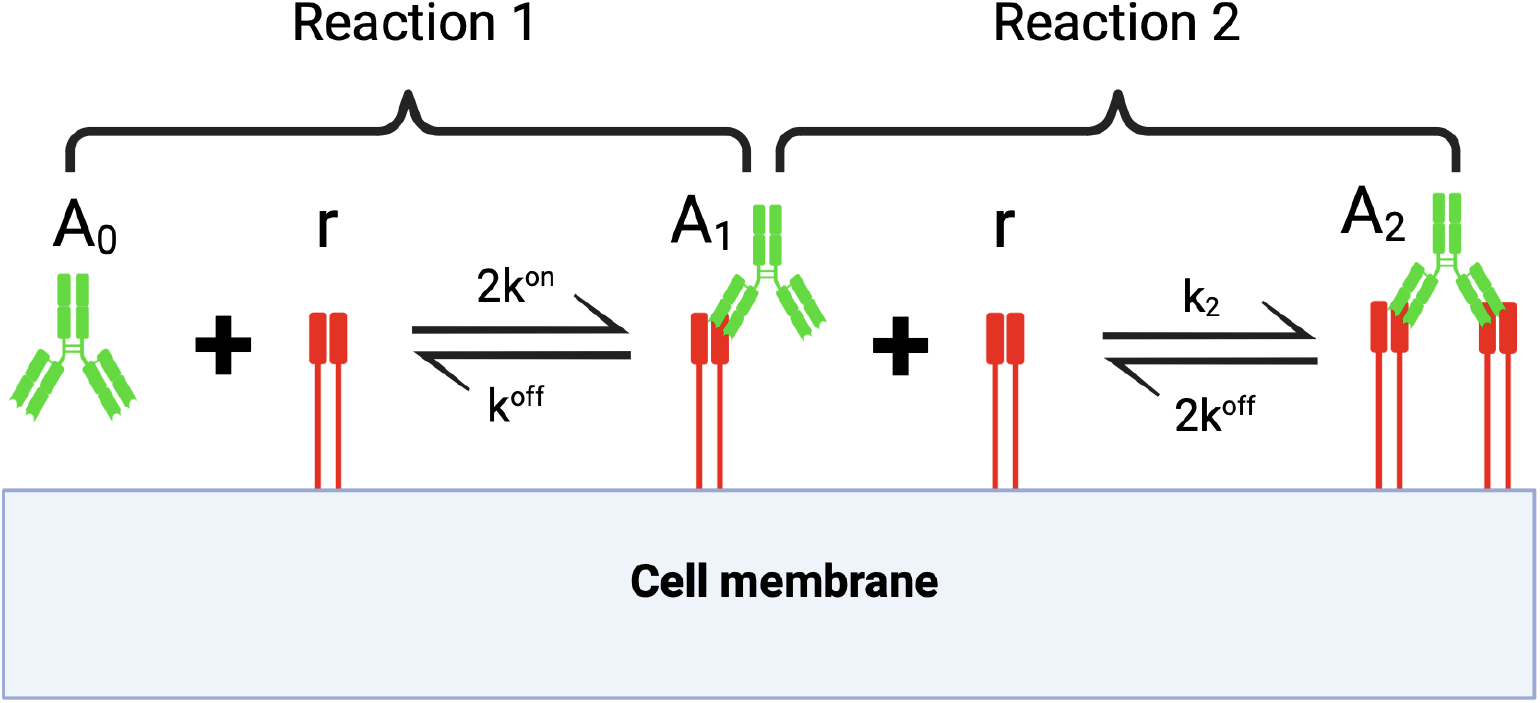
Reaction scheme for antibody binding on the surface of a target cell. An unbound antibody, *A*_0_, binds reversibly with a free target antigen, *r*, at a rate *k*^on^, to form a monovalently bound antibody *A*_1_ and can dissociate at a rate *k*^off^. *A*_1_ can bind another free antigen at a rate *k*_2_, to form a bivalently bound antibody *A*_2_. A bivalently bound antibody can dissociate one of its bound arms away from the target antigen independently of, and at the same rate as, the monovalently bound arm. Note, in an abuse of notation, here *A*_0_ denotes a single unbound antibody whereas in Equations (1)-(4) it denotes the total number of unbound antibodies (similarly for all other variables). Created with Biorender.com.

For simplicity, we neglect antigen internalisation. This is justified because, typically, the timescale of antigen internalisation is much slower than the timescale on which antibody binding takes place on the surface of a cell. For example, the internalisation half life of mavrilimumab is approximately 30 minutes to two hours while the timescale of antibody binding can be much less than a second for high antibody concentrations (Birtwistle and Kholodenko 2009; Vainshtein et al. 2014).

Under the above assumptions, our mathematical model can be written as follows

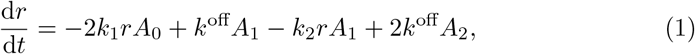

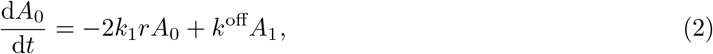

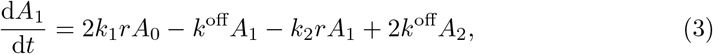

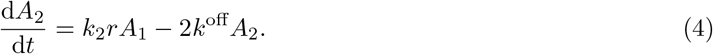

The factor of 2 that appears in the reaction terms 2*k*_1_*rA*_0_ and 2*k*^off^*A*_2_ is due to our modelling assumption that both antibody arms are identical and their dynamics are independent. As a result, the factor of 2 represents when two antibody arms can undertake a reaction (e.g. an antibody is able to bind either of its arms when in solution and, similarly, dissociate either of them when it is bivalently bound). We close Equations (1)-(4) by imposing the following initial conditions:

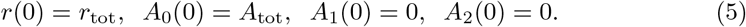

In Equation (5), we assume that all antigens are initially unbound and we denote by *A*_tot_ and *r*_tot_ the total number of antibodies and target antigens respectively within the system. The parameters *A*_tot_ and *r*_tot_ have units of antibody number and antigen number per cell respectively.

By taking suitable linear combinations of Equations (1)-(4), and exploiting Equation (5), it is straightforward to deduce that the total number of antibodies and antigens are each conserved within the system:

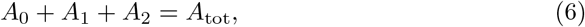

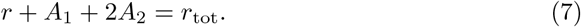

We use Equations (6) and (7) to eliminate *A*_0_ = *A*_tot_ *−A*_1_ *−A*_2_ and *r* = *r*_tot_ *−A*_1_ *−*2*A*_2_ and, henceforth, focus on the following reduced system for *A*_1_(*t*) and *A*_2_(*t*):

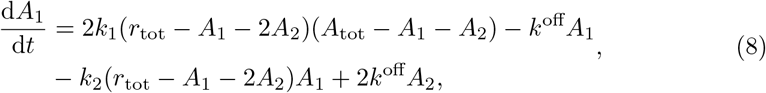

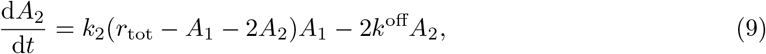

with the following initial conditions:

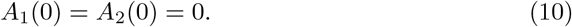

Before analysing our reduced model, we pause to estimate the model parameters.

### 2.1 Model Parameter Estimates

In practice, we model an assay within a well of volume *V*_well_ (units: litres L). Thus, we estimate *A*_tot_, the number of antibodies within the system (with units in numbers of ligand or protein), for a given experimental antibody concentration, by noting that

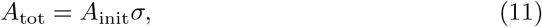

where *A*_init_ is the initial antibody concentration (units: molar concentration M = mol/L), and *σ* a “concentration-to-antibody-number” conversion factor given by

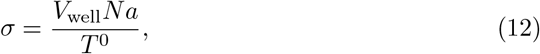

where *Na* = 6.02214 *×* 10^23^ is Avogadro’s number (units: mol^*−*1^) and *T* ^0^ is the target cell number within the assay volume. Equation (12) is normalised with respect to *T* ^0^, because we are focusing on binding to a single target cell.

Within the literature, estimates of *k*^on^ for mAbs typically are stated with units s^*−*1^M^*−*1^. Here, we consider antibody and antigen numbers rather than concentrations. Therefore, we rescale *k*^on^ so its units are consistent with those used in our model:

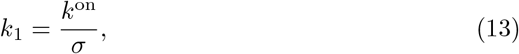

where the units of *k*_1_ are the number of antibodies per second.

Two approaches have been used to model the rate at which the second arm of the antibody binds to cell surface antigens (see reaction 2 in Figure 2). Sengers et al. 2016 assumes the second binding event is limited by antigen diffusion on the cell surface and hence, the reaction rate depends on the diffusion of target antigens. The case of antigens diffusing and binding with antibodies on the surface of a cell is an example of a first-hitting time problem which predicts the time taken for a diffusing body (e.g. an antigen) to encounter a trap (here, an antibody’s binding arm). Following Coombs et al. 2009, the diffusion limited rate constant for the binding of the second arm, *k*_2_, can be estimated to be:

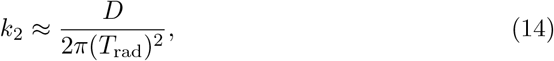

where *D* is the diffusion coefficient of the target antigen (units: m^2^s^*−*1^) and *T*_rad_ is the radius of the target cell (units: metres).

Alternatively, Rhoden et al. 2016 assumes the antigens are immobile and, hence, that the reaction rate is limited by the concentration of free antigens within reach of the bound arm of the antibody. In this case, *k*_2_, the rate of binding of the second arm, is estimated to be:

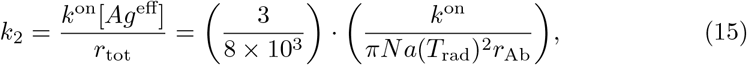

where *k*^on^ was introduced above, [*Ag*^eff^] is the concentration of free antigen within reach of the bound arm of the antibody and *r*_Ab_ is the arm-to-arm binding distance of the antibody.(We refer the interested reader to to Appendix A for more details about the derivation of Equations (14) and (15)).

From here on, we follow Sengers et al. 2016 and assume that the second binding event is driven by antigen diffusion on the cell surface. We justify this choice as follows:

- In Equation (14), the parameter that most commonly varies between target antigens or antibodies is the diffusion coefficient, *D* only. The corresponding parameters in Equation (15) are *k*^on^ and *r*_Ab_. Since *k*^on^ is proportional to the monovalent reaction rate, *k*_1_, (see Equation (13)), the only parameter related to antibody-target interactions that can alter the ratio of *k*_1_ to *k*_2_ is *r*_Ab_. In what follows, we will analyse model outputs as the magnitudes of monovalent and bivalent reactions rates vary. In this vein, the antigen diffusion coefficient, *D*, typically ranges between 10^*−*15^ *−*10^*−*13^ m^2^s^*−*1^ (McCloskey and Poo 1986) but *r*_Ab_, the antibody arm-to-arm distance for an IgG antibody, ranges between 12 *−* 13 *×* 10^*−*9^ m (Sosnick et al. 1992). Therefore, by varying *D* we explore a wider range of values of the bivalent reaction rate *k*_2_.
- In Rhoden et al. 2016, antigens are assumed to be within reach of the antibody binding arm. Assuming that antigens are uniformly distributed over the cell surface, the area swept out by an antibody with *r*_Ab_ = 12.5 *×* 10^*−*9^ m, as a fraction of the surface area of a tumour cell with an average radius of 8 *×* 10^*−*6^ m (Zhou et al. 2019) is approximately 6 *×* 10^*−*7^. Therefore, we estimate that approximately 6 *×* 10^7^ antigens per cell would be needed for one antigen to be within reach of an antibody binding arm. This estimate is much larger than the average number of target antigens on the surface of a tumour cell (10^4^ *−* 10^6^) (Mazor, Oganesyan, et al. 2015). We therefore argue that the rate limiting step for the surface bound reaction is the ability of the target antigen to diffuse until it is within reach of the antibody binding arm, as described by Sengers et al. 2016.

To summarise, we extract values for parameters such as *k*_on_, *k*_off_, *D* and *A*_init_ from the literature and use these to obtain values of *k*_1_, *k*_2_ and *A*_tot_ for use in our model. For reference, the model parameters and their interpretations are provided in Table 1.

**Table 1:**
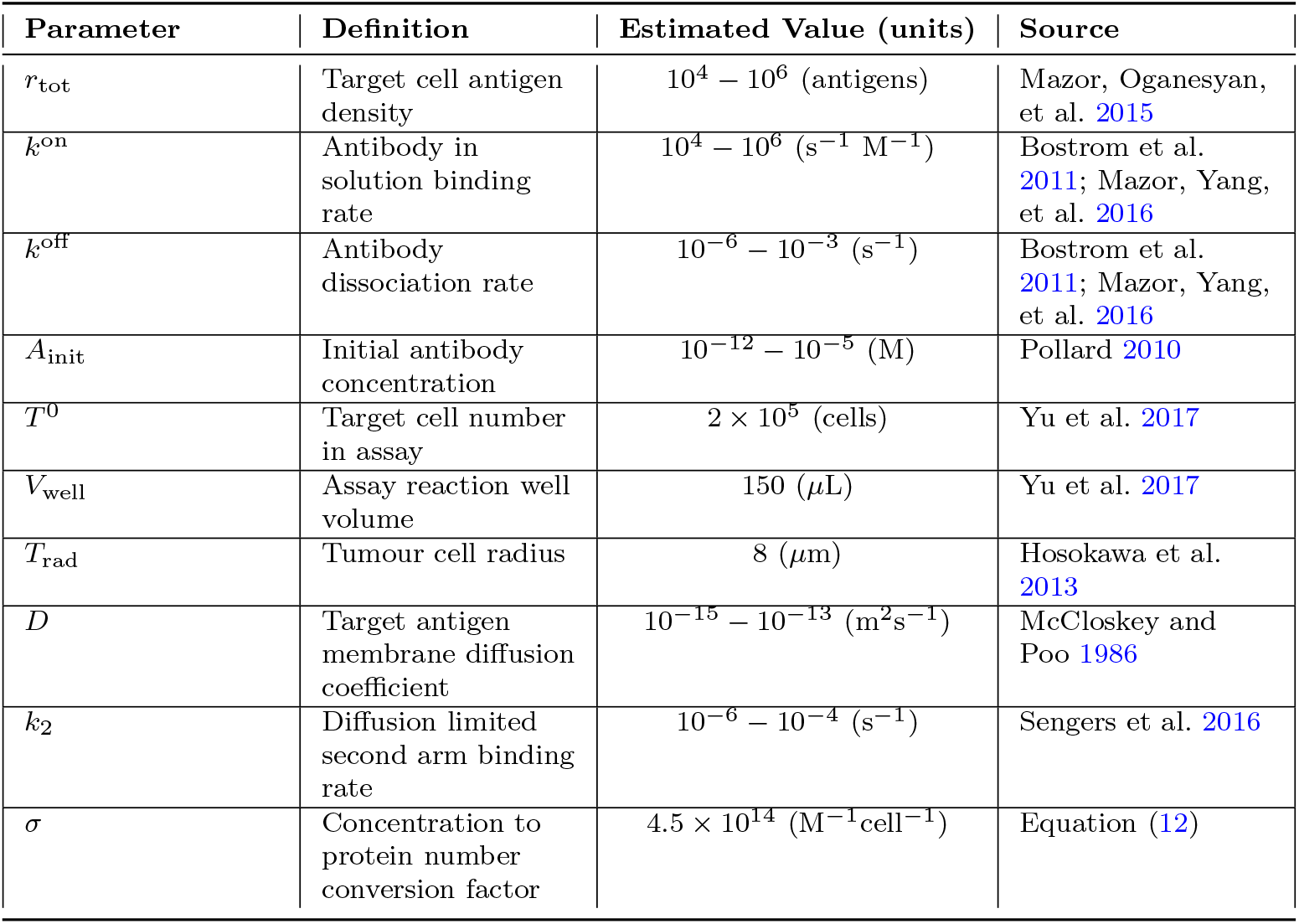
Model parameters associated with Equations (1)-(4). Parameters are from standard human cancer cell lines such as SK-OV3 and mAbs targeting cancer such as trastuzumab (references in table)

### 2.2 Equilibrium Solutions

We now derive equilibrium solutions of Equations (8) and (9). We focus on equilibrium solutions because, for most antibody concentrations, the transient timescale of antibody binding is much shorter (typically less than a second) than the timescales on which effector functions occur (typically minutes to hours) (James and Tawfik 2005; Hoffman et al. 2018). As such, the values of the quantities relating to antibody-antigen binding (e.g. antigen occupancy and the number of bound antibodies) will quickly reach a steady state and we neglect binding dynamics here.

The steady state solutions are determined by setting time derivatives equal to zero in Equations (8) and (9). We denote steady state solutions as *A*_1_ and *A*_2_ by 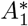 and 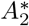 respectively and define 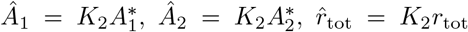 and *Â*_init_ = *K*_2_*A*_init_*σ* where *K*_1_ = *k*_1_*/k*^off^ and *K*_2_ = *k*_2_*/k*^off^. Exploiting these identities and setting d*Â*_2_*/*d*t* = 0 in Equation (9), we have that

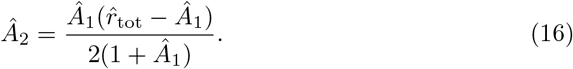

Substituting for *Â*_2_ from Equation (16) into Equation (8), d*Â*_1_*/*d*t* = 0, upon simplification, we obtain the following cubic polynomial for *Â*_1_:

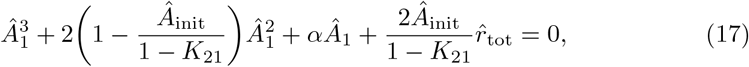

where

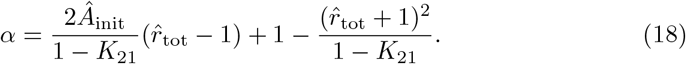

Here, we have introduced *K*_21_ = *K*_2_*/K*_1_ and substituted *A*_tot_ = *A*_init_*σ* from Equation (11) because in what follows we vary the antibody concentration, *A*_init_. It remains to identify the number of real positive roots of Equation (17). From Descartes’ rule of signs, if there is exactly one sign change between the coefficients of a polynomial (where the coefficients follow the order of the power of the polynomial variable), then that polynomial has exactly one real positive root (Anderson et al. 1998). Noting that *K*_21_ *>* 1 because the surface bound reaction rate, *k*_2_, is assumed to be larger than the monovalent reaction rate, *k*_1_, it is possible to show Equation (17) has exactly one real positive root. As d*Â*_1_*/*d*t >* 0 for *A*_1_ = 0 in Equation (8), the trivial solution is unstable and the one real positive root of Equation (17) is the long time asymptote of Equation (8). In subsequent sections, we numerically solve Equation (17), using SciPy’s *fsolve* function to obtain the positive root of Equation (17) (Virtanen et al. 2020).

In subsequent sections, we are particularly interested in the equilibrium values of the following quantities:

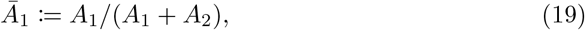

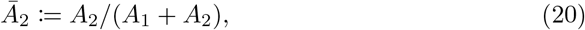

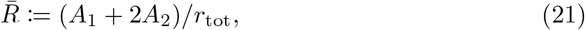

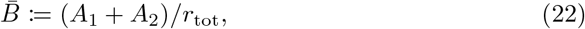

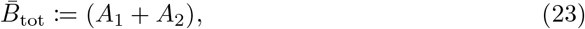

where *Ā*_1_ and *Ā*_2_ are the monovalently and bivalently bound fractions respectively, 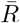 is antigen occupancy, 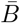 is the bound antibody to antigen ratio and 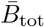 is the total bound antibody number. We are particularly interested in antigen occupancy and bound antibody numbers because they directly impact the effector function of mAbs. For example, the ratio of bound antibody to antigen has been shown to correlate with effector function potency and efficacy (Mazor, Yang, et al. 2016).

### 2.3 Global Sensitivity Analysis Method: Sobol Sensitivity Indices

In this section, we describe Sobol’s method for global parameter sensitivity analysis, prior to its use in Section 3.2 (Sobol′ 2001). Sobol’s method decomposes the variance from a scalar model output into sensitivity indices that quantify the contributions to the variance of different model parameters. Using this method, we can establish which changes in antibody-antigen interactions have the biggest impact in our quantities of interest (e.g. antigen occupancy).

Sobol’s method utilises a quasi-random sequence rather than a uniformly distributed sequence of random numbers to improve convergence of the Sobol sensitivity indices (see Homma and Saltelli 1996 and Sobol′ 2001).

Given a vector of model inputs, ***X*** = *{X*_1_, *X*_2_, …, *X*_*n*_*}*, any model output can be viewed as a function *Y* = *f* (***X***). The variance of the model output, Var(*Y*), can be decomposed as follows

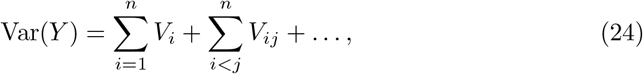

where

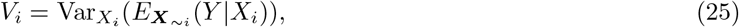

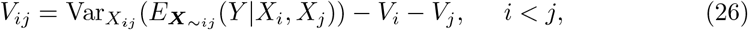

with analogous formulae for higher order terms. Here, ***X***_*∼i*_ is the set of all variables except *X*_*i*_ and ***X***_*∼ij*_ excludes both ***X***_*i*_ and ***X***_*j*_. Equation (24) details how the total variance in model output can be decomposed into variances in the model output given changes in one parameter *X*_*i*_, with *V*_*i*_, and *V*_*ij*_ denoting the variances due to changes in both *X*_*i*_, *X*_*j*_ with iteration to changes in three or more parameters.

A direct measure of the sensitivity of the model output to a single parameter, *X*_*i*_, is called the “first order sensitivity index” or “main effect index” *S*_*i*_ where

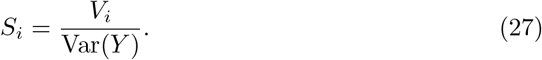

This index measures the effect on the model output *Y* of varying parameter, *X*_*i*_, alone, averaged over variation in all other input parameters. Second order indices, *S*_*ij*_ can be defined in a similar way:

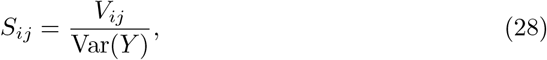

with analogous expressions for higher order interactions. With *S*_*i*_, *S*_*ij*_ and higher order terms defined by Equations (27) and (28), Equation (24) supplies

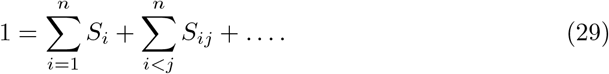

In particular, the sum of first order and higher order indices sum to 1. The indices detailed above describe how the variance in model output depends on a specific parameter. The “Total-effect index” or “Total order index”, *S*_*T i*_, groups together first and higher order indices to measure how the parameter *X*_*i*_ and its interactions with all other parameters affect model output. The total order index, *S*_*T i*_, is given by

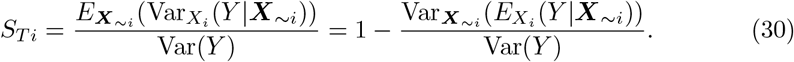

Unlike first and higher order sensitivity indices, the sum over all total order indices is larger than or equal to 1. This is because interactions between parameters are accounted for in each parameters’ total order indices (i.e, the interaction between *X*_*i*_ and *X*_*j*_ is accounted for in both *S*_*T i*_ and *S*_*T j*_), so some terms are counted more than once.

To calculate the Sobol indices, we use the following procedure:

1. Fix upper and lower bounds for the model parameters of interest.
2. Generate multiple parameter sets by sampling from these parameter ranges using the the quasi-random sequence defined in Homma and Saltelli 1996; Sobol′ 2001.
3. For each parameter set generated at step two, calculate the model outputs of interest.
4. For each model output generated at step three, calculate the first and total order sensitivity indices using Equations (27) and (30).

We use the Python package SALib to calculate the sensitivity indices (Herman and Usher 2017; Iwanaga et al. 2022). We also include a dummy variable in our analysis (Marino et al. 2008). The dummy variable does not appear in the model and is used as a threshold, to identify significant parameters above artefact in the sensitivity analysis. In particular, the model’s outputs should not be sensitive to the dummy variable.

## 3 Results

### 3.1 Analysis of Antibody-Target Interactions

In this section, we analyse the equilibrium solutions predicted by the model to establish which parameters influence the quantities defined by Equations (19)-(22) in different experimental systems, noting that antigen expression levels and antibody numbers can vary over many orders of magnitude. To capture this, we define the ratio of total antibody to antigen number in the system as

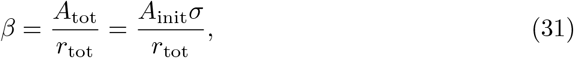

where the constant is defined by Equation (12). Of interest is how the quantities in Equations (19)-(22) change as *β* varies. We will investigate the total bound antibody, 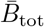 in the next section. In practice, the value of *β* can range over many orders of magnitude; when *β ≪* 1, antigens are in excess of antibody and similarly, when *β ≫* 1, antibodies are in excess of antigens.

In Figure 3, we show how the equilibrium solutions of Equations (16) and (17) and related quantities of interest change as *β* varies. Regarding the fractions of monovalently and bivalently bound antibodies, *Ā*_1_ and *Ā*_2_, Figure 3 shows that when the value of *β* is low (10^*−*1^ *−* 10^1^), most antibodies are bivalently bound and there are few monovalently bound antibodies. As the value of *β* increases (10^2^ *−* 10^6^), antibodies switch from being bivalently to monovalently bound.

**Fig. 3:**
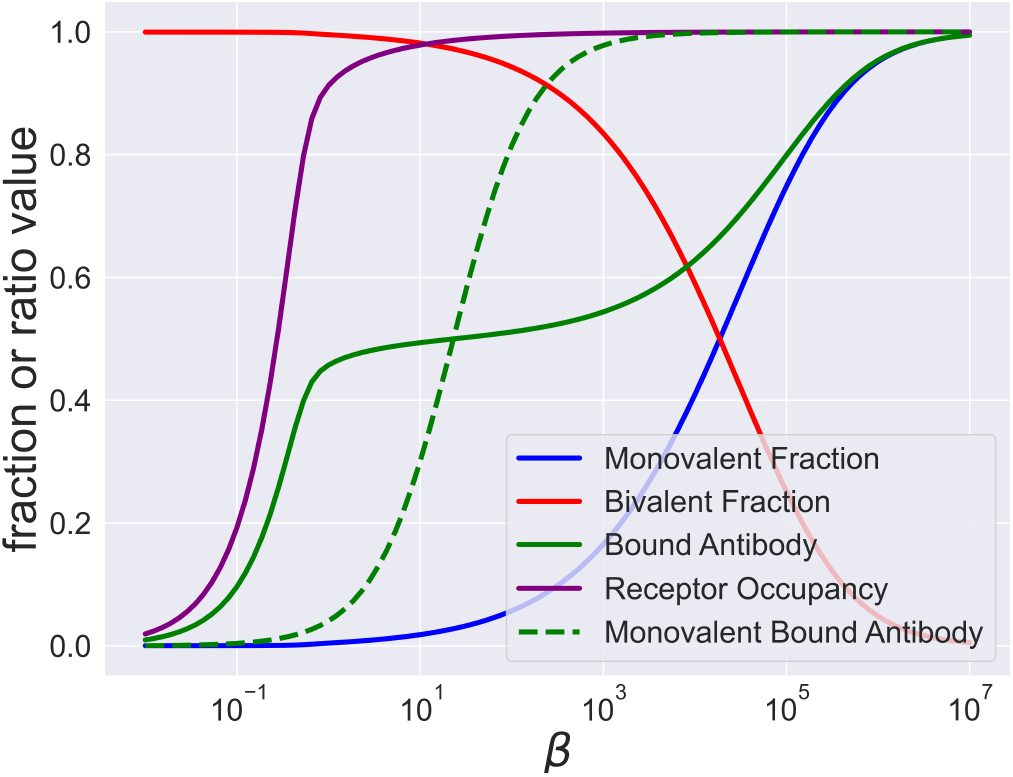
Series of curves showing how the equilibrium values of fraction of monovalently bound antibodies ((*Ā*_1_= *A*_1_/(*A*_1_ + *A*_2_, blue curve), fraction of bivalently bound antibodies (*Ā*_2_= *A*_2_/(*A*_1_ + *A*_2_), red curve), antigen occupancy(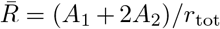), purple curve) and the bound antibody to total antigen number ratio (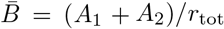, green curve) change as *β* varies. For comparison, 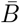 is plotted for the case of a monovalent antibody which is modelled by setting *k*_2_ = 0 and *A*_2_(0) = 0 (green dashed curve. Values of *A*_1_ and *A*_2_ were obtained by solving Equations (16) and (17) with *k*^on^ = 10^5^ s^*−*1^M^*−*1^, *k*^off^ = 10^*−*4^ s^*−*1^ and *D* = 10^*−*14^ m^2^s^*−*1^.

Figure 3 shows further that 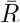 approaches one (i.e. antigens are saturated) for small values of *β* (*β ≈* 𝒪 (10^1^)). This is because antibodies each bind two antigens so few antibodies are required to saturate the target antigens. From Figure 3, we also see that antigens remain saturated as antibodies transition from being bivalently to monovalently bound, while the ratio of bound antibody to total antigen level, 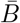, changes. In particular, 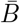 reaches a plateau when *β ≈* 𝒪 (10^1^), at which antibodies are primarily bivalently bound, and then increases as the number of monovalently bound antibodies increases. For comparison, in Figure 3 we also plot 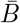 for a monovalent antibody (green dashed line). As expected, monovalent antibodies achieve maximum numbers of bound antibody at a lower value of *β* than bivalent antibodies. This is consistent with the observations of Mazor, Yang, et al. 2016 that monovalent antibodies increase the number of antibodies bound to the cell surface and elicit enhanced ADCC potency. A much larger value of *β* (*β ≈* 𝒪(10^7^)) is required for a bivalent antibody to achieve the same number of bound antibodies as a monovalent antibody.

Given that a high value of *β* may correspond to either a large number of antibodies or a small number of antigens, it is unclear how the quantities defined in Equations (19)-(22) depend on each of *A*_init_ and *r*_tot_. In Figure 4 we calculate *Ā*_1_, *Ā*_2_, 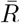 and 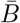 as *A*_init_ and *r*_tot_ vary. Figures 4a and 4b show that there are distinct favourable regimes for the monovalently and bivalently bound fractions, *Ā*_1_ and *Ā*_2_; *Ā*_1_ attains its highest value when the value of *r*_tot_ is low and *A*_init_ is high (high value of *β*). Furthermore, there are more bivalently than monovalently bound antibodies (*Ā*_2_ > *Ā*_1_ for most values in the (*r*_tot_, *A*_init_) plane while the antibody concentration that maximises 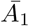 is inversely proportional to the value of *r*_tot_. Even for the smallest values of *r*_tot_ however, a large value of *A*_init_ is required to maximise *Ā*_1_.

**Fig. 4:**
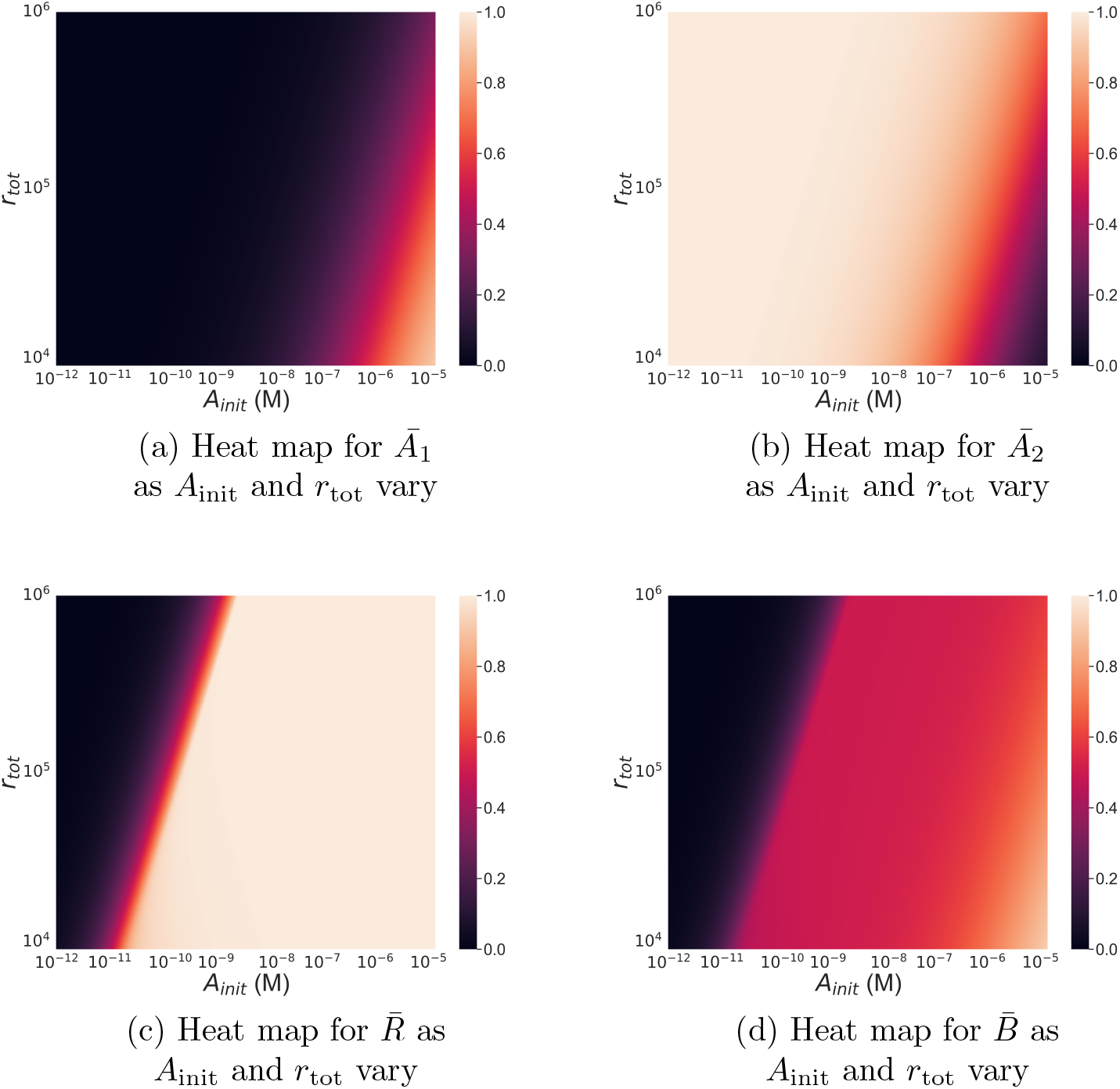
Heat maps showing how (a): the equilibrium fraction of monovalently bound antibodies, Ā_1_ = *A*_1_/(*A*_1_ + *A*_2_), (b): fraction of bivalently bound antibodies, Ā_2_ = *A*_2_/(*A*_1_ + *A*_2_), (c): antigen occupancy, 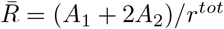 and (d): bound antibody to total antigen number ratio 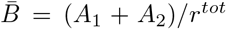 change as the parameters *A*_init_ and *r*_tot_ vary. Values of *A*_1_ and *A*_2_ were obtained from Equations (16) and (17) with *k*^on^ = 10^5^ s^*−*1^M^*−*1^, *k*^off^ = 10^*−*4^ s^*−*1^ and *D* = 10^*−*14^ m^2^s^*−*1^.

The results in Figures 3, 4a and 4b are consistent with the experimental work of Bondza et al. 2020. Our results predict that antibodies are primarily bivalently bound for most antibody concentrations and monovalently bound only for very high concentrations. Similarly, Bondza et al. 2020 found that the binding state of rituximab and obinutuzumab changed in a dose-dependent manner, with the antibody primarily bivalently bound for most antibody concentrations and monovalently bound only for very high concentrations. The results in Figures 4a and 4b extend those of Bondza et al. 2020, by predicting that the dose-dependent binding behaviour they observed does not depend on the antibody concentration alone. Instead, this behaviour depends on *β*, the ratio of total antibody to antigen within the system with antibodies becoming primarily monovalently bound when the value of *β* is high (large *A*_tot_, small *r*_tot_). In particular, antibodies may be primarily bivalently bound even for high antibody concentrations if the value of *β* is not large enough. This can be seen in Figures 4a and 4b when the value of *r*_tot_ is high (*r*_tot_ = 10^6^) and the value of *β* is smaller.

To assess how the behaviours observed in Figures 4a and 4b translate to quantities that relate to effector function potency and efficacy, in Figures 4c and 4d we plot equilibrium values of antigen occupancy, 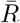, and the ratio of bound antibody to total antigen number, 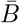. There is a curve in the (*r*_tot_, *A*_init_) plane that separates regions in which antigens are saturated 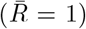 from those where antigens are unsaturated 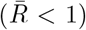. In fact, *β ≈* 1*/*2 on this curve as this value of *β* gives the minimum number of antibodies required to saturate the target antigens (the value is 1*/*2 because each antibody can bind two antigens). Therefore, for smaller values of *r*_tot_, a smaller value of *A*_init_ is required to give *β* = 1*/*2. This is to be expected as smaller numbers of target antigens require fewer antibodies to saturate them. This corresponds to a “tight binding regime” where the binding affinity is larger than the available antigen concentration and, as a result, the reaction is limited by the small number of antigens (Jarmoskaite et al. 2020).

As in Figure 3, Figure 4d shows that 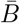 attains its maximum value at the same values of *r*_tot_ and *A*_init_ as 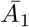. Naturally, the ratio of bound antibody to total antigen number increases with the number of monovalently bound antibodies because more antibodies can bind to the cell surface if they are only binding a single target antigen. Mazor, Yang, et al. 2016 showed how binding affinity can be adjusted to maximise the number of bound antibodies (see Figure 1). Our analysis extends the results of Mazor, Yang, et al. 2016 by predicting which total antigen numbers and antibody concentrations increase the number of monovalently bound antibodies (and as a result the number of bound antibodies) leading to increased effector function potency and efficacy. In particular, our model predicts the largest improvement in effector function when the value of *β* is high (e.g. large value of *A*_init_ and low value of *r*_tot_).

### 3.2 Global Sensitivity Analysis for Antigen Occupancy and Number of Bound Antibodies

In this section, we conduct a global parameter sensitivity analysis (Iooss and Lemaıtre 2015) for a range of antibody concentrations to examine the dependence of key model outputs on model parameters (details of how we calculate the sensitivity indices are included in Section 2.3 (Sobol′ 2001)). The outputs of interest are the equilibrium values of 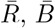 and 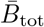 (see Equations (21)-(23)).

As before, we focus on these quantities as they have been shown to correlate with the potency and efficacy of mAb treatments (Mazor, Yang, et al. 2016). We utilise Sobol’s method to perform the global sensitivity analysis, and examine the sensitivity of 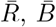 and 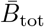 to variations in *k*^on^, *k*^off^, *r*_tot_ and *D*. We choose these parameters because *k*^on^ and *k*^off^ are commonly reported for mAbs while *r*_tot_ and *D* can vary depending on the target antigen. When calculating the sensitivity indices, we vary *k*^on^, *k*^off^, *r*_tot_ and *D* over the ranges in Table 1.

In Figure 5 we present total order sensitivity indices for 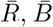 and 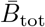 We calculate these indices for a series of fixed values of the antibody dose *A*_init_. Focusing on antigen occupancy, 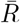, (Figure 5a), we observe two qualitatively different regimes. For small values of *A*_init_ (10^*−*11^ *−* 10^*−*9^ M), the sensitivity is dominated by the antigen density, *r*_tot_. For larger values of *A*_init_ (10^*−*8^ and 10^*−*5^ M), 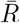 is more sensitive to the binding rates.

**Fig. 5:**
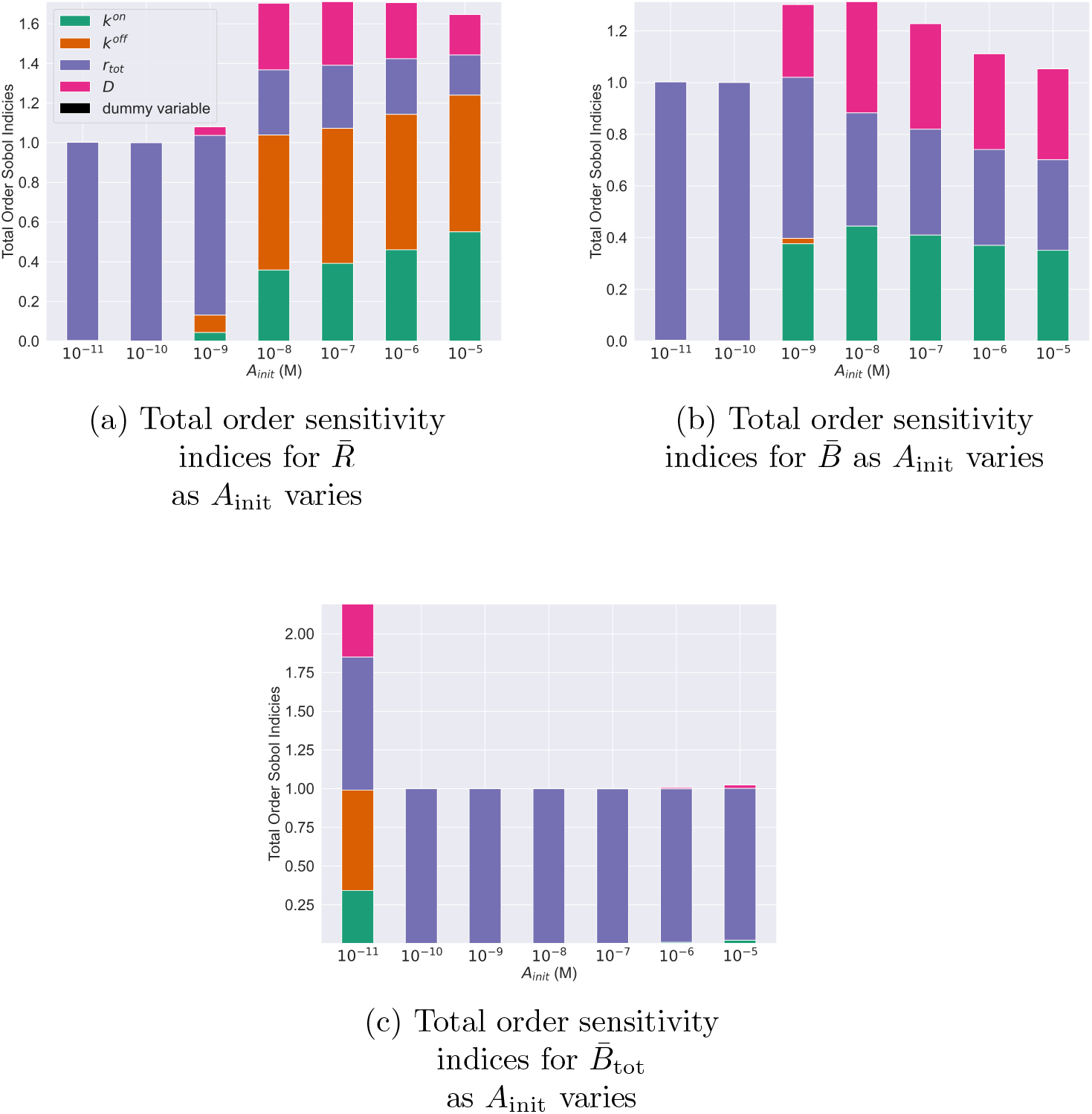
Total order Sobol sensitivity analysis at different fixed values of *A*_init_ for (a) antigen occupancy 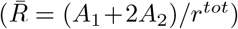, (b) bound antibody to total antigen number ratio 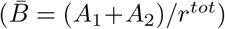, and (c) total number of bound antibodies 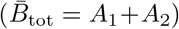. A dummy parameter was added to the analysis to estimate uncertainty within the sensitivity indices as in Marino et al. 2008.

We observe a similar dependence of the sensitivities of the ratio of bound antibody to total antigen number, 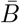, on the antibody concentration (Figure 5b). For small values of *A*_init_ (10^*−*11^ *−* 10^*−*9^ M) the sensitivity is dominated by the antigen density. As *A*_init_ increases, 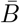 becomes sensitive to the binding parameters. Interestingly, comparing Figures 5a and 5b shows that 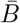 is less sensitive to the off rate, *k*^off^, than 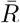 is.

Figure 5c shows that the total number of bound antibodies, 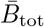 is most sensitive to the total antigen number, *r*_tot_, for all values of *A*_init_ except *A*_init_ = 10^*−*11^ M. This behaviour is expected since the maximum number of antibodies that can bind to a cell is bounded by the total number of target binding sites when antibodies are in excess. The reason 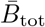 is sensitive to the binding parameters when *A*_init_ = 10^*−*11^ M is predicted to be because antigens are excess and, therefore, the number of bound antibodies is limited by the antibody’s ability to bind antigens rather than to the total antigen number.

### 3.3 Investigation of the Avidity Effect

In this section, we use the model to investigate the avidity effect for antibody binding. Recall that the avidity effect is described as the apparent increase in binding affinity due to multiple bindings. Antibodies can elicit an avidity effect by binding antigens with more than one of their antigen binding arms.

One way to quantify EC50, the binding potency of an antibody, is to measure the antibody concentration at which half of the maximum binding signal is obtained. We measure the avidity effect, ΔEC50, by comparing EC50 values for a monovalent (EC50_Monovalent_) and a bivalent (EC50_Bivalent_) antibody (see Figure 6). We define ΔEC50 as

**Fig. 6:**
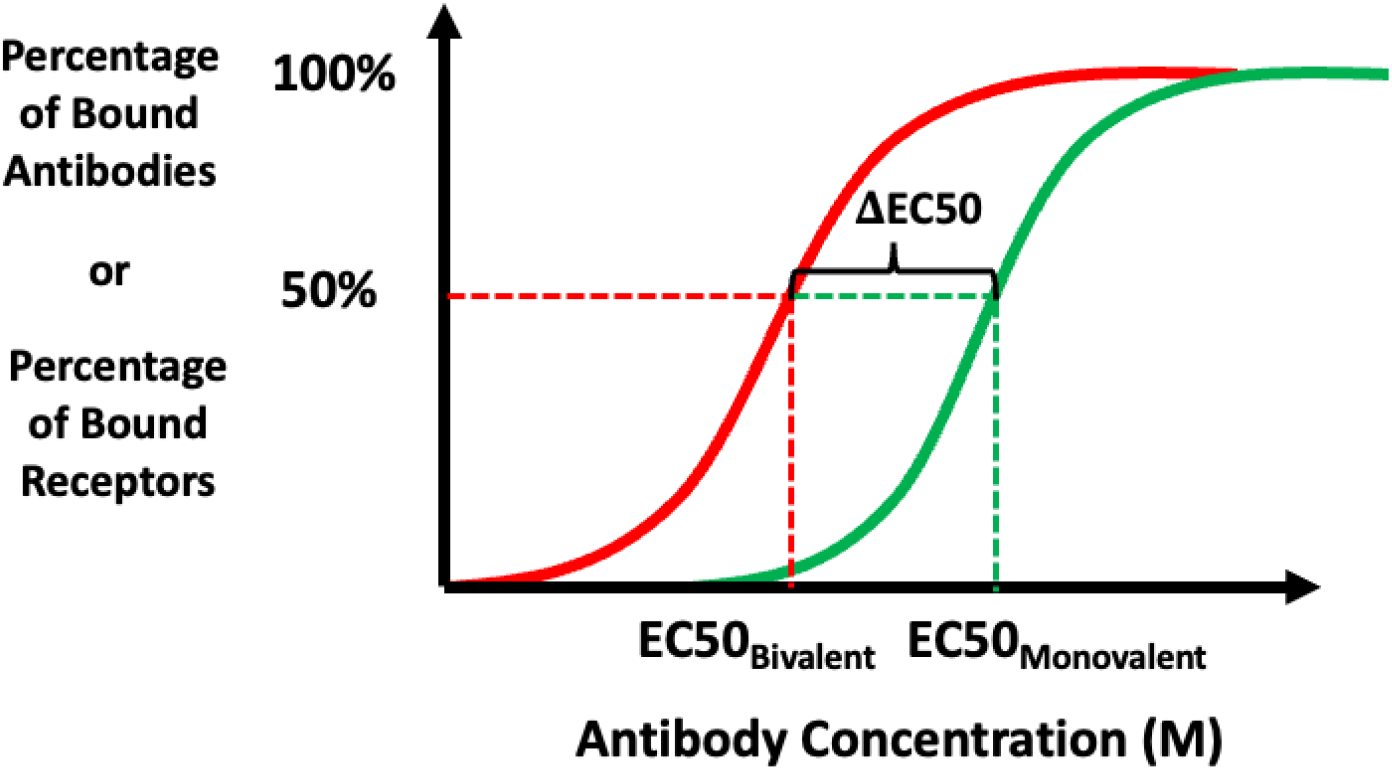
A schematic showing the avidity effect is measured with ΔEC50. The EC50 value is defined as the antibody concentration at which the measurement is half of its maximum value. Here, EC50_Bivalent_ and EC50_Monovalent_ are the iEC50 values for a bivalent and corresponding monovalent antibody (i.e, an antibody with only one functional arm, so that *k*_2_ = 0, *A*_2_ = 0 in Equations (8) and (9)). All other parameters are kept the same between the two curves. The shift in EC50, termed ΔEC50, between the monovalent and bivalent case is due to multiple arms binding antigens, i.e, the avidity effect.

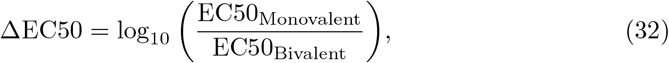

where EC50_Bivalent_ is calculated using Equations (8) and (9), with EC50_Monovalent_ calculated from the monovalent analogue of these equations, obtained by setting *k*_2_ = 0 and *A*_2_(0) = 0. We quantify ΔEC50 in two different ways; by considering the binding signal to be the number of bound antibodies, 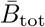, or to be antigen occupancy, 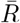.

In Figure 7, we show how ΔEC50 changes as we vary the target antigen density, *r*_tot_, diffusion coefficient, *D*, and binding affinity as measured by the dissociation constant, *K*_*D*_ = *k*^off^*/k*^on^. Focusing on the avidity effect for 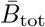, in Figures 7a, 7c and 7e, there are regions in the (*r*_tot_, *K*_*D*_) plane where the value of ΔEC50 is high. In particular, if the antigen density is high and the antibody does not bind strongly (*r*_tot_ = 10^6^ and *K*_*D*_ = 10^*−*6^ M), then the avidity effect is large (ΔEC50 = 4). In contrast, the avidity effect is smaller when there are few antigens and the antibody binds strongly (*r*_tot_ = 10^3^ and *K*_*D*_ = 10^*−*10^ M). The region of parameter space in which the avidity effect is large (ΔEC50 *≥* 2) increases with *D* because as the diffusion coefficient increases the second antibody arm binds more readily. As a result, we observe a strong avidity effect in Figure 7e compared to Figures 7a and 7c, even when the antibody is a strong binder (*K*_*D*_ = 10^*−*10^ M).

**Fig. 7:**
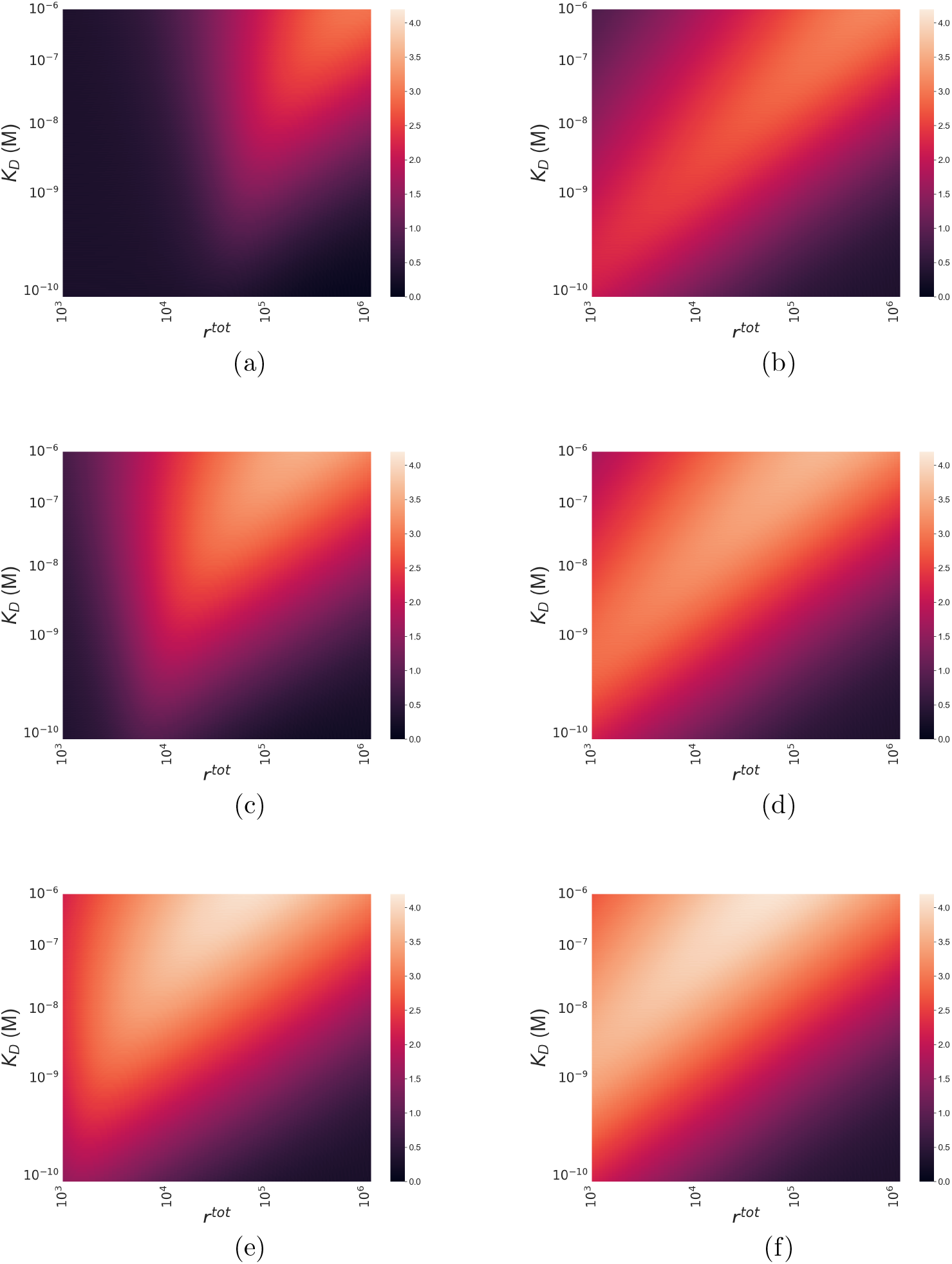
Heat maps using Equation (32) showing how the value of ΔEC50 changes with *r*_tot_ and *K*_*D*_ for different values of *D*. ΔEC50 was calculated for the number of bound antibodies 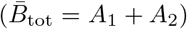 *A*_2_) and 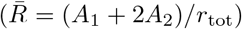. (a): ΔEC50 for 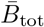 with *D* = 10^*−*15^ m^2^s^*−*1^. (b): ΔEC50 for 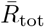 with *D* = 10^*−*15^ m^2^s^*−*1^. (c): ΔEC50 for 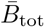 with *D* = 10^*−*14^ m^2^s^*−*1^. (d): ΔEC50 for 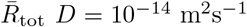. (e): ΔEC50 for 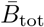 with *D* = 10^*−*13^ m^2^s^*−*1^. (f): ΔEC50 for 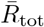 with *D* = 10^*−*13^ m^2^s^*−*1^.

Inspection of Figures 7b, 7d and 7f also reveals that there are values of *r*_tot_ and *K*_*D*_ for which the avidity effect for 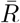 is enhanced. With the exception of *D* = 10^*−*13^ m^2^s^*−*1^, these regions are markedly different to those for 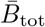 (Figure 7a, 7c and 7e). In particular, the avidity effect for 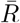 is large when the antibody is a strong binder (low value of *K*_*D*_) for all reported values of *D* shown. However for small values of *K*_*D*_, a large avidity effect for 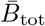 is only seen when *D* = 10^*−*13^ m^2^s^*−*1^ (Figure 7e).

Figure 7 suggests that the avidity effect for 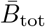 is large when bivalent binding is favoured. Mazor, Yang, et al. 2016 showed that lowering an antibody’s binding affinity favours monovalent binding while Figure 4 shows that a large antibody to antigen ratio also increases monovalent binding. This explains why the avidity effect is small when *r*_tot_ and *K*_*D*_ are small and large when *r*_tot_ is high; these conditions favour monovalent and bivalent binding respectively.

## 4 Discussion and Conclusions

Monoclonal antibodies are an important class of immmunotherapies that are being used to treat cancer. Central to their mechanism of action are their interactions with their target antigens. During the drug discovery process, the binding affinity between target and antibody is often prioritised. However, other factors, such as the antigen density and valency of the antibody, are known to be important for effector function potency and efficacy (Mazor, Yang, et al. 2016).

In this paper, we have used a mathematical model of a bivalent, monospecific antibody binding to cell membrane antigens given by Equations (1)-(4), to investigate how changing model parameters impacts antigen occupancy, the number of bound antibodies and the avidity effect. We focused on these quantities as they have been shown to relate to the potency and efficacy of antibody effector functions such as ADCC (Mazor, Yang, et al. 2016). We used the model to investigate how *β* = *A*_tot_*/r*_tot_, the ratio of the number of antibodies to antigens within the system, impacts antigen occupancy, the ratio of bound antibody to antigen and whether the antibody is monovalently or bivalently bound (Figures 3 and 4). We found that higher values of *β* resulted in larger numbers of monovalently bound antibodies and, consequently, an increase in the number of antibodies bound to the cell surface. This result suggests that a possible way to increase ADCC efficacy is to ensure that the antibody dose is large enough that monovalent binding is favoured. In turn, increased numbers of monovalently bound antibody will increase the number of antibody Fc regions available to activate an effector cell to lyse a tumour cell

Next, we performed a global parameter sensitivity analysis to determine how quantities that relate to mAb potency and efficacy, such as antigen occupancy, change when parameters that govern antibody-target interactions (on and off rates *k*_on_ and *k*_off_, surface diffusion of antigen *D*, and total antigen number *r*_tot_) are varied. We performed the sensitivity analysis for different fixed values of the antibody concentration, *A*_init_, to see how the sensitivities changed with antibody dose and as a result, *β*. We identified two regimes of sensitivity for antigen occupancy: a low *β* regime where antigens are in excess of antibody and antigen occupancy is most sensitive to antigen density, and a high *β* regime where antibody is in excess of antigens and antigen occupancy becomes sensitive to the binding parameters. Both regimes may be inferred from Figure 5a, where for low values of the antibody concentration, *A*_*init*_ (*A*_*init*_ *≈* 10^*−*11^ *−* 10^*−*9^, low *β* regime) the total order sensitivity index for the total antigen number is large, with a value of approximately one. As the initial antibody concentration increases (*A*_*init*_ *≥* 10^*−*8^ M) the total order sensitivity indices for *k*^on^ and *k*^off^ become large, attaining values of approximately 0.4 and 0.6 respectively.

The sensitivity analysis for the bound antibody to total antigen number ratio also shows two regimes depending on whether the value of *β* is high or low (Figure 5b). Similar to antigen occupancy, the sensitivities to the binding parameters increase at higher antibody concentrations (*A*_*init*_ *≥* 10^*−*9^ M). Interestingly, in the high *β* regime, the sensitivity to *k*^off^ of the ratio of bound antibody to total antigen number is negligible whereas antigen occupancy is sensitive to *k*^off^ at the same antibody concentrations.

The dependence of the sensitivities on antibody concentration has important implications for the development of antibody therapeutics. If the tumour microenvironment, or other factors such as antibody pharmacokinetics, hinder antibody infiltration then the ratio of antibody to antigen will be low (low *β* regime). In such situations, targets of high abundance should be prioritised and increasing the affinity of the antibody may not significantly enhance the mAb’s anti-tumour effects. Conversely, if antibodies are in surplus (high *β* regime), then altering the binding affinity is predicted to impact antigen occupancy and the bound antibody to total antigen number ratio, potentially improving mAb potency. This is the case for immune checkpoint inhibitors, that benefit from increased antigen occupancy, and effector functions, such as ADCC, that benefit from maximising the number of bound antibodies per antigen.

Our analysis predicts that there are (at least) two ways to increase the potency and efficacy of effector functions that relate to the number of bound antibodies. Figure 5b shows that the choice of parameters to be varied in order to maximise potency and efficacy depends on the antibody concentration. In particular, Figure 5b indicates a lack of sensitivity to the binding parameters at low antibody concentrations. Therefore, altering the binding affinity to improve effector function potency and efficacy, as implemented by Mazor, Yang, et al. 2016, is predicted to have no effect for these concentrations. Only when the antibody concentration is sufficiently large (e.g. *A*_init_ *≥* 10^*−*8^ M) will altering the affinity potentially improve effector function. Additionally, Figure 5c highlights that increasing the total antigen number will have the greatest effect on the total number of bound antibodies.

The above observations suggest ways to increase the number of effector cell activating antibody Fc regions on the surface of a tumour cell. While increasing the ratio of bound antibody to antigen will maximise the number of antibody Fc regions on the surface of a tumour cell for a given total antigen density, increasing the total antigen number will increase the upper bound for the number of antibody Fc regions. The results presented here suggest that utilising both strategies will be advantageous for mAb effector functions. In particular, choosing targets that are overly expressed on tumour cells is already common practice and will increase the maximum number of antibody Fc regions that can potentially be presented to the effector cell. Ensuring the antibody to antigen ratio is high and altering the binding affinity to enhance monovalent binding, as in Mazor, Yang, et al. 2016, will maximise monovalent binding and ensure that the number of antibody Fc regions on the surface of the tumour cell approaches its maximum given by the total antigen number.

Another focus of this work has been to study the avidity effect seen in antibody binding. We have shown that there exists ranges of the total antigen number, *r*_tot_ and the binding affinity, as measured with *K*_*D*_ = *k*^off^*/k*^on^, where the avidity effect is predicted to significantly increase the EC50 for both antigen occupancy and the number of bound antibodies. Figure 7 shows that the avidity effect is predicted to generate a significant change in EC50 for different ranges of target antigen numbers and binding affinities depending on whether the antigen occupancy or the number of bound antibodies is measured. In particular, the size of the avidity effect is small for the number of bound antibodies compared to antigen occupancy when there are few target antigens (low value of *r*_tot_) and the affinity for the target is high (low value of *K*_*D*_). It follows that, in this situation, an antibody whose anti-tumour effects correlate with the number of bound antibodies is predicted to gain little benefit from the avidity effect. In contrast, an antibody whose potency and efficacy relies on antigen occupancy, such as immune checkpoint inhibitors, should gain an improvement in their anti-tumour effects. These results can be used to aid target selection in preclinical development of therapies that utilise the avidity effect.

In summary, we have used mathematical modelling to study the interaction of mAbs with their antigens in order to understand how this may regulate mAb effector functions in cancer immunotherapies, noting that factors affecting the number of bound antibodies on a target cell, antigen occupancy and the avidity effect can impact the potency and efficacy of the mAb therapy. With prospective relevance for the preclinical development of immuno-oncological therapies, the main results of this study are threefold. Firstly we have predicted the importance of the ratio of antibody to antigen within the system, *β*, on the antibody’s binding state and the values of this ratio that enhance antigen occupancy and bound antibody numbers, expanding on previous experimental work by Mazor, Yang, et al. 2016 and Bondza et al. 2020. In addition, we have identified parameter sensitivities of measures of mAb potency and efficacy and predict that these sensitivities can be dependent on antibody concentration. Finally, we have highlighted regions of parameter space that are predicted to possess a large avidity effect, with parameters associated with large avidity differing between two measures of binding signal, the number of bound antibodies and antigen occupancy.

## Bullet Point Summary

### What is already known

- Antibody-target interactions such as the avidity effect are key to mAb potency and efficacy.
- Binding affinity can be adjusted to improve antibody effector function potency and efficacy.

### What does this study add

- The key antibody-antigen interactions and antibody concentrations that are predicted to impact mAb potency and efficacy.
- For what parameter ranges a large avidity effect is predicted.

### What is the clinical significance

- We predict what conditions are necessary to improve mAb treatment potency and efficacy and the avidity effect.

## Funding

This work was supported by funding from the Engineering and Physical Sciences Research Council (EPSRC) [grant number EP/S024093/1]. For the purpose of Open Access, the author has has applied a CC BY public copyright licence to any Author Accepted Manuscript (AAM) version arising from this submission.

## Conflict of Interest Statement

The authors declare no conflict of interest.

**Appendix A Modelling Methodologies for the**

**Second Binding Event**

In this appendix, we look in more detail at modelling the binding of the second arm, as described in Sengers et al. 2016 and Rhoden et al. 2016.

## A.1 Method 1: Surface Diffusion Limited Binding

In Sengers et al. 2016, an ODE model is developed to describe bispecific antibody binding to two different targets, CD4 and CD70, on the surface of a target cell. For an antibody bound to a cell with one arm and an additional free antigen to bind, their separation on the cell surface must be on the molecular scale. Sengers et al. 2016 assume that, due to this small binding distance, the reaction where the antibody binds its second arm with a target antigen on the surface of the cell is independent of the in-solution binding rate. Instead, they suppose that the rate limiting step is reactant diffusion i.e. the ability of the reactants diffusing on the cell surface and get within binding distance of each other. In particular, the rate at which the antibody is able to bind a second antigen on the cell surface is assumed to be proportional to the sum of the diffusion coefficients of the reactants.

Sengers et al. 2016 also estimate that the timescale on which a antigen can freely diffuse across large distance across the cell surface is much smaller than the average half-life of an antibody-antigen complex. As such, a second diffusion-driven binding event is possible even when antigens are sparsely distributed across the cell surface.

For antigens diffusing on the surface of a cell and binding with each other, this is an example of a first hit time problem which is extensively studied in Coombs et al. 2009. From Coombs et al. 2009, we have that *k*_2_, the rate of binding of the second arm is

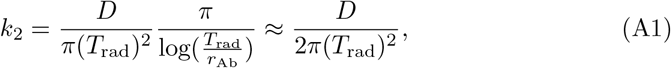

where *D* is the diffusion coefficient of the target antigen (units: m^2^s^*−*1^), *T*_rad_ = 8*×*10^*−*6^ is the average radius of a tumour cell (units: m) and *r*_Ab_ = 1.25 *×* 10^*−*8^ is the arm-to-arm antibody length (units: m). For most cell surface proteins, *D* lies in the range 10^*−*15^ *−* 10^*−*13^ m^2^s^*−*1^ which gives estimates of *k*_2_ in the range of 10^*−*6^ *−* 10^*−*4^ (units: s^*−*1^).

## A.2 Method 2: Free Antigen Concentration Within Antibody Arm Reach

Rhoden et al. 2016 use an ODE model to investigate the binding kinetics of a bispecific antibody as target antigen numbers and binding affinities vary. In their model, the binding of the bispecific antibody’s arms are viewed as independent events. When modelling the binding of the second arm, Rhoden et al. 2016 assume that the antibody is fixed to a region near the cell surface and the unbound arm of the antibody is confined to a volume defined by its size, flexibility and structure (see Figure A1). In more detail, the antibody samples a hemisphere of radius equal to the length of a typical IgG. The concentration of target antigens within the hemispherical volume is estimated from data recording the average number of antigens per cell.

The effective target concentration, [*Ag*_*eff*_], after the first binding event is estimated using the following formula

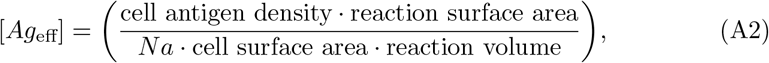

or, equivalently,

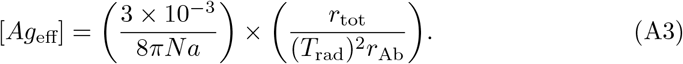

Here, *r*_tot_ is the number of target antigens per cell, *T*_rad_ and *r*_Ab_ are the radius of the target cell and arm-to-arm antibody length respectively (units: m). The factor 10^3^ converts metres cubed (m^3^) to litres (L) so [*Ag*_eff_] has units of molar concentration.

**Fig. A1:**
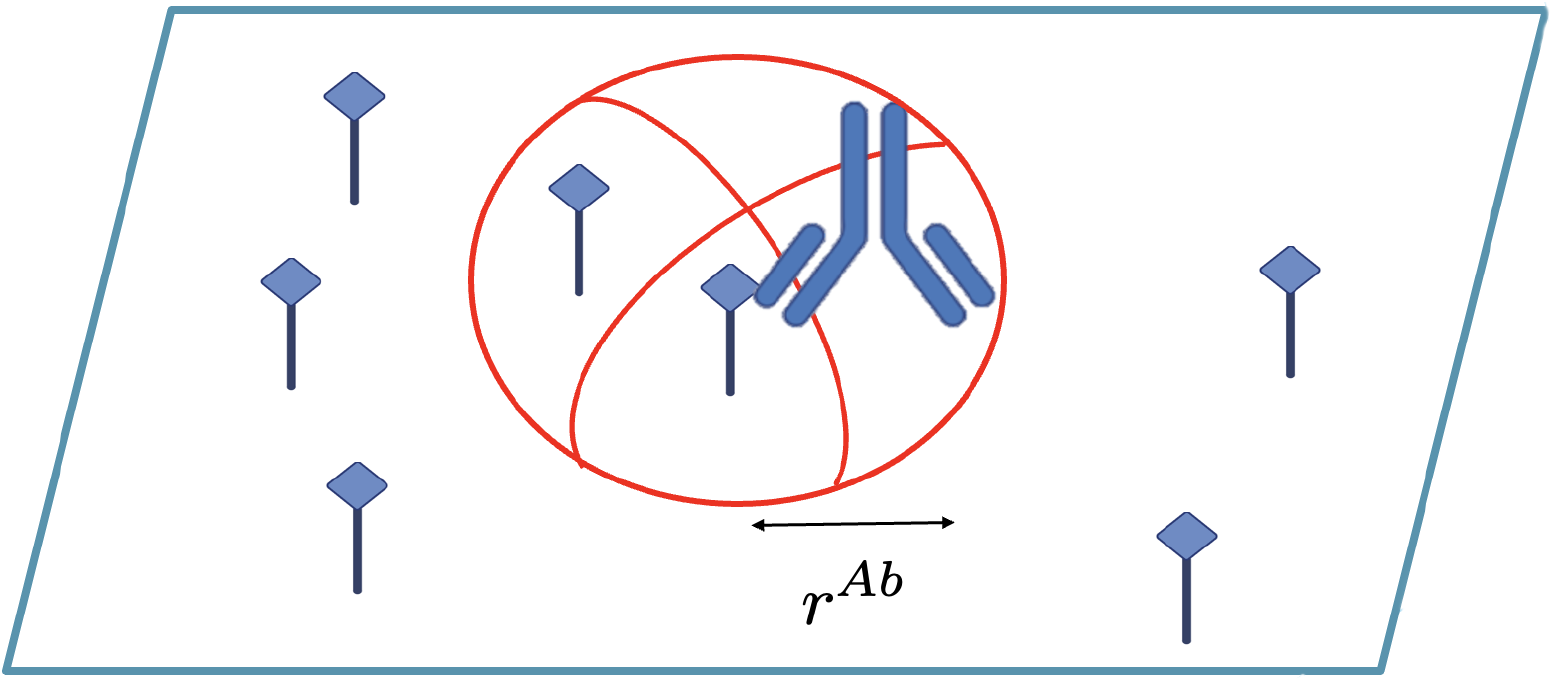
Depiction of the constrained volume that the antibody can sample from after the first binding event. *r*_Ab_ denotes the arm to arm distance of a standard IgG antibody.

The rate constant for the second binding event, *k*_2_, is then

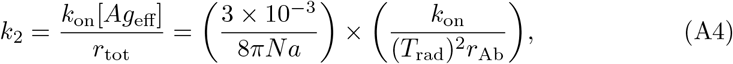

where *k*^on^ is the in-solution monovalent binding reaction rate (units: s^*−*1^M^*−*1^). In Equation (A4), we divide by *r*_tot_ so that *k*_2_ is a reaction rate per antibody-antigen complex. This step was not necessary in Equation (A1) as the diffusion-limited rate describes a single antibody-antigen complex reaction.

## Notes

### Competing Interest Statement

The authors have declared no competing interest.

